# Genomewide association study reveals transient loci underlying the genetic architecture of biomass accumulation under cold stress in Sorghum

**DOI:** 10.1101/760025

**Authors:** Nadia Shakoor, Erica Agnew, Greg Ziegler, Scott Lee, César Lizárraga, Noah Fahlgren, Ivan Baxter, Todd C. Mockler

## Abstract

*Sorghum bicolor* is a promising cellulosic feedstock crop for bioenergy because of its potential for high biomass yields. However, in its early growth phases, sorghum is sensitive to cold stress, preventing early planting in temperate environments. Cold temperature adaptability is vital for the successful cultivation of both bioenergy and grain sorghum at higher latitudes and elevations, and for early season planting or to extend the growing season. Identification of genes and alleles that enhance biomass accumulation of sorghum grown under early cold stress would enable the development of improved bioenergy sorghum through breeding or genetic engineering. We conducted image-based phenotyping on 369 accessions from the sorghum Bioenergy Association Panel (BAP) in a controlled environment with early cold treatment. The BAP is a collection of densely genotyped and racially, geographically, and phenotypically diverse accessions. The plants were weighed, watered, and imaged daily to measure growth dynamics and water use efficiency (WUE). Daily, non-destructive imaging allowed for a temporal analysis of growth-related traits in response to cold stress. We performed a genome-wide association study (GWAS) to identify candidate genomic intervals and genes controlling response to early cold stress. GWAS identified transient quantitative trait loci (QTL) strongly associated with each growth-related trait, permitting an investigation into the genetic basis of cold stress response at different stages of development. The analysis identified *a priori* and novel candidate genes associated with growth-related traits and the temporal response to cold stress.

**SIGNIFICANCE STATEMENT:** Genome-wide association study of bioenergy sorghum accessions phenotyped under early season cold stress revealed transient QTLs for highly heritable biomass and growth-related traits that appeared as the temperature increased and plants developed. Sorghum accessions clustered into multiple groups for each heritable trait with distinct growth profiles. GWAS identified candidate genes associated with growth traits and cold stress responses. The top-performing accessions with the highest growth-related trait values over time and temperature shifts will be useful for further genetic analysis and breeding or engineering efforts directed at biomass yield enhancements.

## INTRODUCTION

*Sorghum bicolor* (L.) Moench, a C_4_ crop native to Africa and known for its drought and heat tolerance, is a promising bioenergy crop because of its ability to grow in marginal environments and a high potential for biomass yield (1). Early season planting, which extends the growing season of bioenergy sorghum and potentially increases exposure to rainfall is a strategy to maximize biomass yield (2, 3). Cultivation in northern regions and at higher elevations could increase the land available for bioenergy sorghum production, without using land areas needed for food or feed crop production.

However, early planting and cultivation in temperate areas will not lead to higher biomass yields if plant development stalls. Because of its cold-sensitivity, planting sorghum in temperatures below 12-15°C will diminish the yield (4, 5, 6). Therefore, identification of sorghum accessions with enhanced tolerance to cold is needed to lengthen the growing season, expand growing regions, and to achieve higher total biomass yields.

Increased cold tolerance is only one of several traits necessary for improved biomass yield. Identification of ‘ideotype-positive’ and ‘ideotype-negative’ accessions can prioritize germplasm with multiple positive characteristics that may contribute to increased yield and other desirable traits through breeding or engineering. For example, accessions with high biomass, reduced height, and high WUE may be more desirable for breeding bioenergy sorghum. Alternatively, accessions that are tall but have low biomass and low WUE may be the least desirable for multiple reasons, including a higher propensity for lodging (7). Crosses of lines exhibiting these different ideotypes can also be used to create resources such as nested association mapping (NAM) populations to elucidate the genetic architectures underlying the desired characteristics.

The genetic basis of sorghum’s response to cold stress and its potential for cold adaptability must be better understood in order to breed or engineer sorghum lines that can thrive in the lower temperatures of early spring and in colder climates. Cold tolerance is a complex quantitative trait, and there is phenotypic variability for cold tolerance among sorghum accessions (8, 9). Natural genetic variation in sorghum’s response to cold stress has also been identified (10). Genome-wide association mapping of cold sensitivity traits in diverse germplasm is a promising approach to identify allelic variation that may be harnessed to improve the cold tolerance of sorghum.

The aim of this study was to analyze the phenotypic variability and genetic architecture of bioenergy sorghum’s response to early cold stress conditions in a controlled environment. We used a high throughput imaging-based system to collect daily phenotypic measurements from a set of 369 diverse accessions from the sorghum Bioenergy Association Panel (BAP) (1) genotyped with 232,303 high-quality single nucleotide polymorphism (SNP) markers (S1). Daily phenotypic measurements are more beneficial than endpoint measurements because the response to cold stress can change over time and as plants develop. Therefore, regular imaging and phenotyping allowed for the comparison of the cold stress response in the BAP at different developmental stages.

High throughput image-based phenotyping was performed to track growth over time and under early cold stress in this sorghum panel. GWAS on the daily phenotypic data revealed genetic variation underlying the response to early season cold stress and identified transient QTLs related to biomass, height, hull area, water use efficiency (WUE) and relative growth rate (RGR). Candidate genes associated with these phenotypes were identified and may prove useful for crop improvement through breeding or targeted genetic modifications.

## RESULTS

### Population Structure and Kinship

In this study, we tested 369 accessions of the BAP grown under early cold stress conditions. The BAP (1) comprises six subpopulations that represent a racial, geographic, and phenotypically diverse selection of sorghum accessions. The BAP was designed to limit variation in key bioenergy traits, such as height and sensitivity to photoperiod, within a range of desirable values (1) (S1). We generated a kinship matrix showing the genetic correlations in the BAP (S2). The kinship matrix revealed distinct groups of highly correlated accessions. In many of the highly correlated clusters, the individual accessions shared characteristics such as the country of origin and similar photoperiod sensitivity. For example, in one cluster, 47 of 48 accessions originated from Ethiopia. Another cluster comprised 29 accessions, and 25 of those originated from Ethiopia, while 26 were photoperiod-sensitive. One cluster was comprised entirely of photoperiod-insensitive accessions, while 29 of 30 accessions in another cluster were of the cellulosic type. These clusters reflect an uneven population distribution and illustrate the necessity for controlling for population structure in this study.

### Germination and Seedling Vigor

Seeds were sown in pots at 15°C and grown at that temperature for 31 days. The temperature was then increased to 24°C for seven days and then increased to 32°C for the final 18 days of the experiment. Plant growth and development were observed using daily, non-destructive imaging throughout the eight-week experiment. Germination dates were determined for each plant by image analysis and manual validation. For 369 accessions, at least two plants germinated. Of these 369 accessions, 41 germinated within ten days after planting (DAP) during the 15°C cold treatment, and the remaining accessions germinated between 11 DAP and 46 DAP (S3). Accessions that failed to germinate or died by 46 DAP were excluded, leaving 309 accessions for subsequent phenotypic and genetic analyses (S3).

### Trait Heritability and Phenotypic Variation

To analyze the phenotypic response to early cold stress in the BAP, we evaluated five bioenergy-related traits of interest: biomass, height, hull area, water use efficiency (WUE), and relative growth rate (RGR). Images captured daily from three angles and the amount of water each plant received daily enabled calculation of phenotypic measurements for each plant for each day of the experiment.

We used the three replicates for each accession to calculate broad-sense heritability within the experiment for each of the traits at each DAP to determine the proportion of phenotypic variation explained by genetic variation in the BAP. The heritability for each trait changed over time and as the temperature increased (Fig. 1a), generally increasing through the early growth period and decreasing as the plants became larger and more root bound. Heritability estimates ranged from 0.544 to 0 (S4). Height at 44-54 DAP was the most heritable trait with a heritability range of 0.544 to 0.503. The heritability values were moderate (0.493 to 0.3) for height at 23-43 DAP, height at 55-56 DAP, WUE at 36-45 DAP, and hull area at 42-56 DAP. The remaining traits had low heritability (0.297-0).

**Figure 1.**
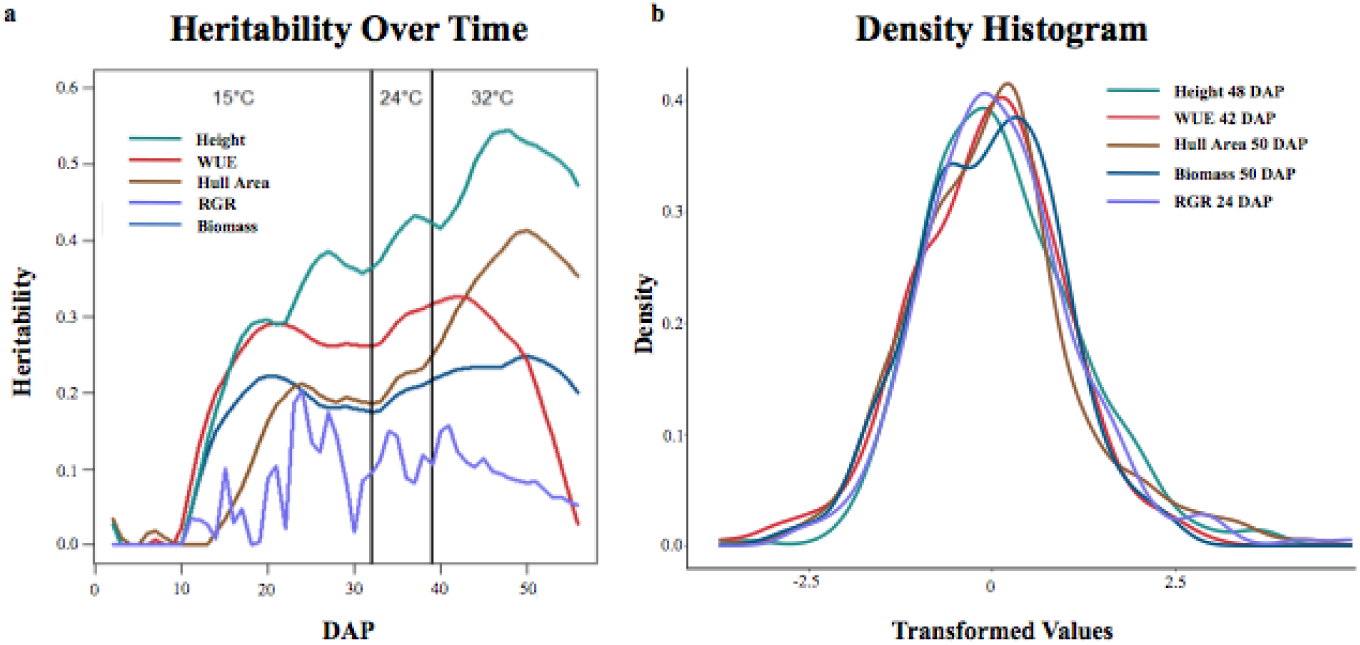
Traits of interest show distinct trends in heritability over time and temperature changes. (a) Heritability of traits over time at each temperature (15°C, 24°C, and 32°C). (b) Density histograms of each trait analyzed show the day at which each corresponding trait is most heritable (height at 48 DAP, WUE at 42 DAP, hull area at 50 DAP, biomass at 50 DAP, and RGR at 24 DAP). Each trait was transformed into an approximately normal distribution using the Box-Cox method.

Heritability of WUE increased between 10-20 DAP, remained relatively flat until 42 DAP, then decreased. The decreased heritability of WUE was expected and coincides with an increased uncertainty of biomass calculations and water use estimates during the final two weeks of the experiment. There was a significant increase in the overlapping of leaves after 42 DAP, resulting in the underestimation of plant biomass from the imaging data. The heritability for RGR had no discernable trend. The range of variation observed for the phenotypes of interest is depicted in the density histogram (Fig. 1b). The density histogram shows the frequency of the mean of the analyzed phenotypes at the DAP at which each trait had the highest heritability.

### Trait Correlations

The correlations among biomass, height, hull area, RGR, and WUE were evaluated at 51 DAP (Fig. 2). The plants began to overlap and grow outside of the camera view after that date, making area measurements based on pixel numbers less indicative of the actual plant size. Biomass was most highly correlated to WUE with a r^2^ of 0.95 but also showed high correlations to hull area and height. The correlation between biomass and RGR was negative, with a r^2^ of -0.52.

**Figure 2.**
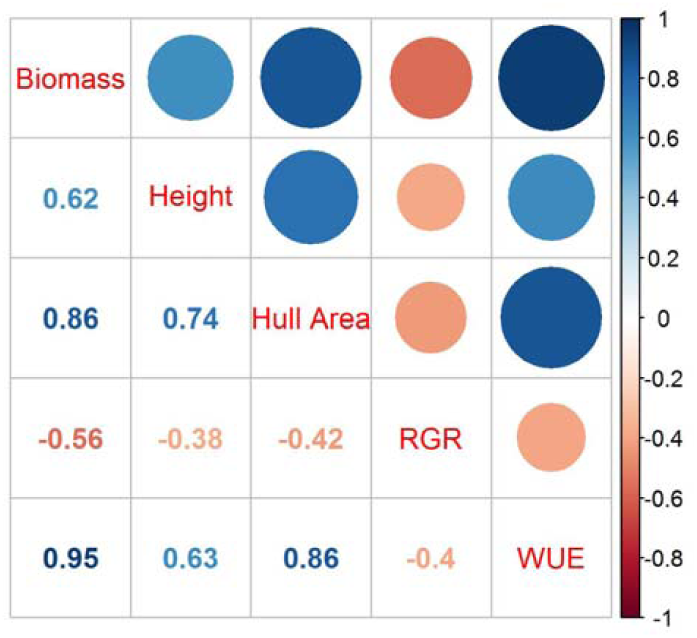
Correlations Among Traits at 51 DAP. Correlations among biomass, height, hull area, RGR, and WUE at 51 DAP. Dark blue represents the strongest correlation, and dark orange represents the weakest correlation. The r^2^ for each correlation is in the corresponding box below the colored circles.

### Phenotypic Ranking of Accessions

The daily plant images were analyzed to quantify the biomass, hull area, height, and WUE phenotypes for each DAP. Based on the resulting values, for each trait, we ranked the accessions based on their performance. Some accessions were ranked high or low for multiple growth-related traits. The Venn diagram (Fig. 3) illustrates overlaps among the top or bottom 5% of accessions at 51 DAP. Table S5 provides a complete ranking of accessions for each trait at 51 DAP. Five accessions (PI152771, PI55064, PI514456, PI569454, and PI534165) are represented in the bottom-ranked accessions for all four traits. PI154988, PI535794, PI569148, PI535792, PI329584, and PI535793 are among the lowest-ranked accessions for biomass, hull area, and WUE, while PI569457 is among the lowest-ranked accessions for biomass, hull area, and height. Six additional accessions ranked as poor performers for two traits of interest: PI452544, PI552851, PI570038, and PI570254 for biomass and WUE; PI157035 for hull area and height; and PI540518 for hull area and WUE. Among the top-ranked accessions, PI19770, PI329299, PI455301, and PI453106 were in the top 5% for biomass, height, hull area, and WUE. PI329403 and PI452619 were among the top accessions for biomass, hull area, and WUE, while PI92270 was a top performer for hull area, height, and WUE and PI563222 was among the top lines for biomass, height, and WUE. Finally, eight additional lines ranked among the top accessions for two traits.

**Figure 3.**
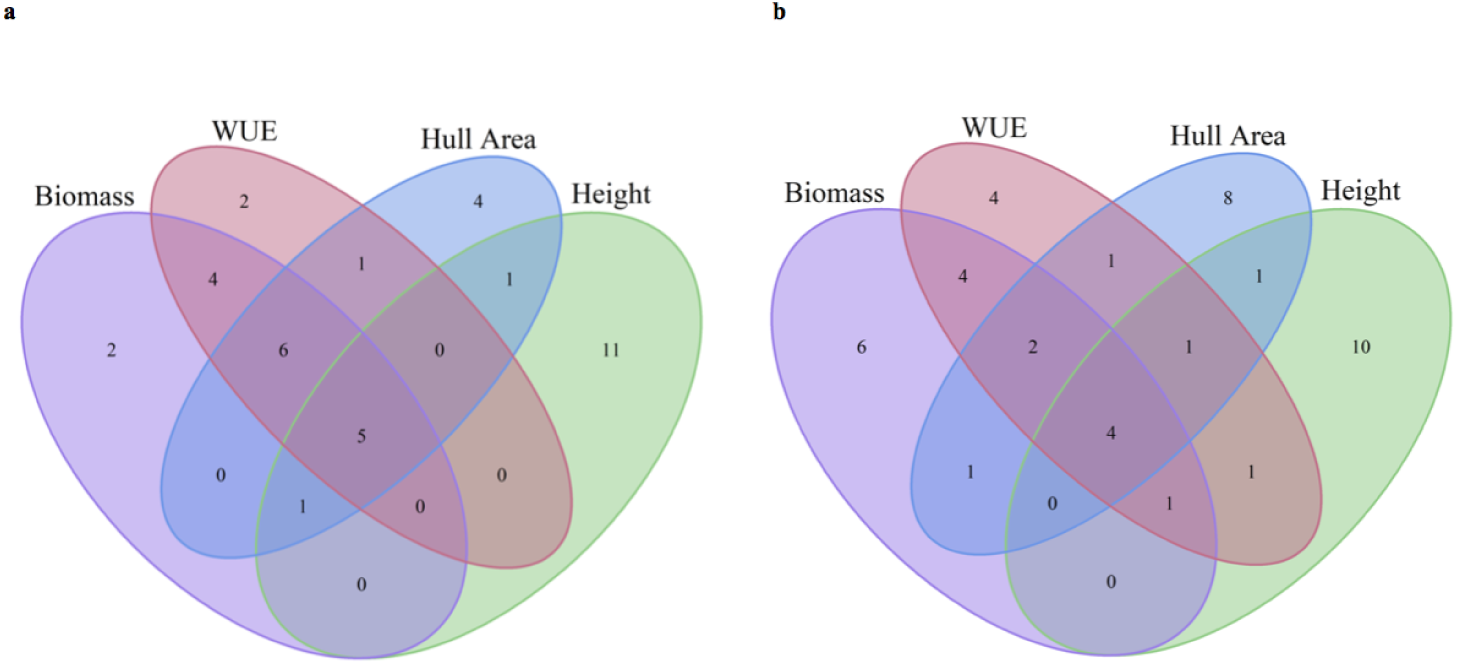
Overlaps of High and Low Performing Accessions at 51 DAP. Venn diagrams depict numbers of accessions represented in the lowest-performing accessions (a) or highest-performing accessions (b) that overlap for biomass, WUE, hull area, or height at 51 DAP.

### Ideotype-Positive and Ideotype-Negative Accessions

Comparisons of accession rankings among multiple traits facilitated the identification of ‘ideotype-positive’ and ‘ideotype-negative’ accessions. Accessions ranked within the top 10% for WUE but with a height-to-biomass ratio in the bottom 5% of accessions were designated as ideotype-positive (Fig. 4a). There were five ideotype-positive accessions in the early period of the experiment, from 15 DAP to 31 DAP at 15°C: PI329286, PI329517, PI329711, PI455221, and PI562730. There were three ideotype-positive accessions in the late period of the experiment, from 39 DAP to 56 DAP after the temperature had been increased to 32°C: PI329471, PI329517, and PI585461. Accession PI329517 was ideotype-positive throughout the experiment.

**Figure 4.**
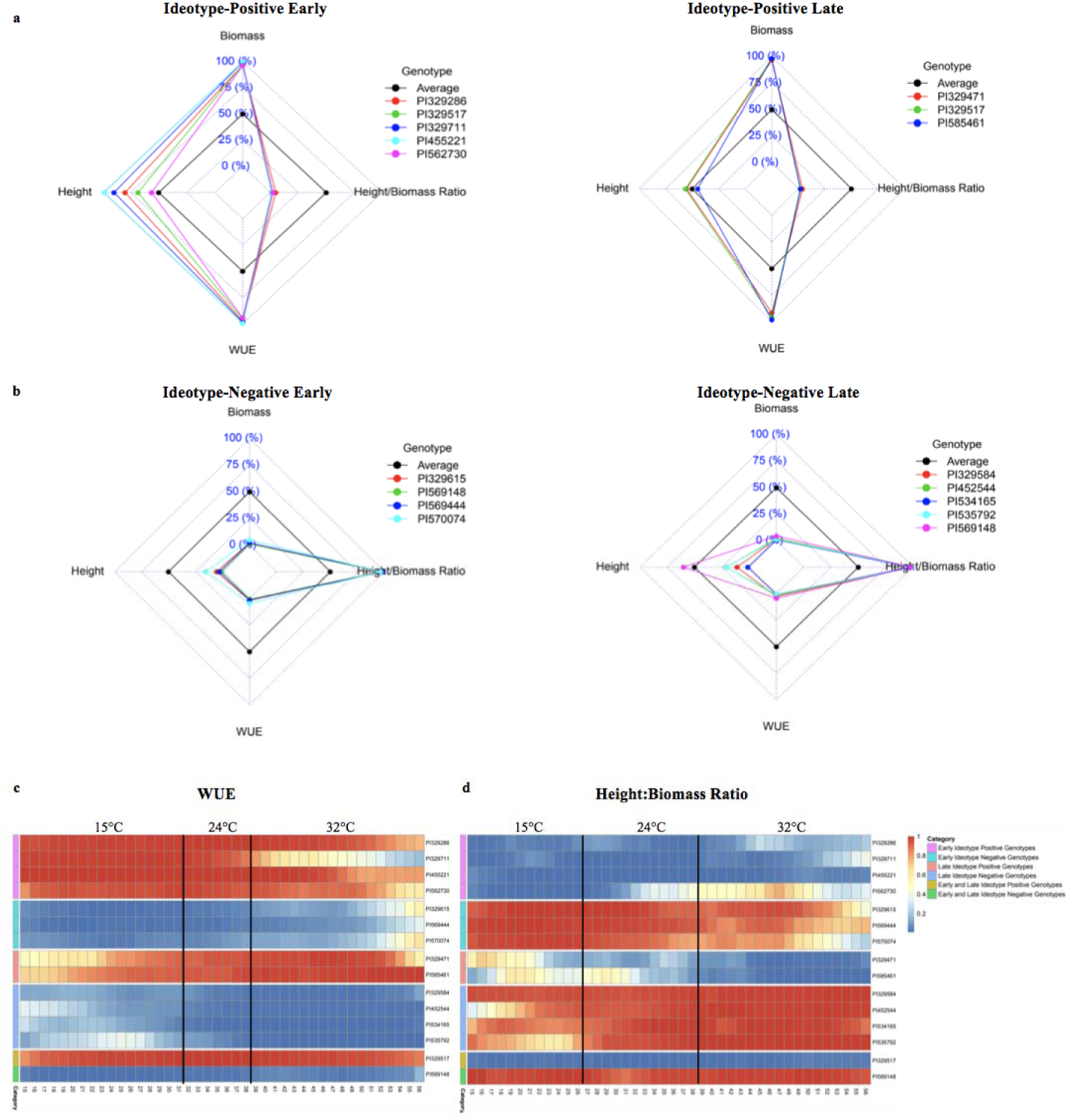
Ideotype-Positive and Ideotype-Negative Accessions for WUE and Biomass-Associated Traits. (a) Early and late ideotype-positive radar plots show accessions within both the top 10% of WUE and lowest 5% of height:biomass ratios. Each dashed square represents a percentile range: 0% is the center-most square, and 100% is the outermost square. Accessions whose phenotypes fall within a given percentile range for a given trait (height, height:biomass ratio, or WUE) reside on the inside of the corresponding square. (b) Early and late ideotype-negative radar plots show accessions within both the lowest 10% of WUE and the highest 5% of height:biomass ratios. (c-d) Heatmaps show the change in rank of early and late ideotype-positive and negative accessions over time and at each temperature stage for WUE (c) and height:biomass ratios (d). Dark blue indicates the lowest WUE (an ideotype-negative trait) and lowest height:biomass ratio (an ideotype-positive trait). Dark orange indicates the highest WUE (an ideotype-positive trait) and highest height:biomass ratio (an ideotype-negative trait).

Accessions within the bottom 10% for WUE accessions but with a height-to-biomass ratio in the top 5% of accessions were designated as ideotype-negative (Fig. 4b). There were four early ideotype-negative accessions: PI329615, PI569444, PI569148, and PI570074, and five late ideotype-negative accessions: PI329584, PI452544, PI534165, PI535792, PI569148. Accession PI569148 maintained an ideotype-negative phenotype throughout both the 15 to 31 DAP and 39 to 56 DAP periods. Heatmaps show the WUE (Fig. 4c) and height-to-biomass ratios (Fig. 4d) of the early and late ideotype-positive and ideotype-negative accessions from 15 DAP through 56 DAP.

### Temporal WUE Profiles and Clustering of Accessions

The daily quantification of traits provided the opportunity to assess temporal changes in trait values as the temperature increased and plants developed. For brevity, we only refer to the five top and bottom-ranked accessions for WUE, a key trait. Table S6 provides the complete ranked list of accessions for each trait. Figure 5 shows the temporal WUE profiles for the five highest- and lowest-ranked accessions at each temperature stage. The top-ranked accessions for WUE at each temperature exhibited a WUE profile over time distinct from the profiles of the accessions ranked lowest for WUE accessions, and the separation became more apparent as the plants developed and the temperature increased (Fig. 5). Even at 15°C, the highest-ranked accessions for WUE are distinguishable with a steeper increase in WUE over time compared to the bottom-ranked accessions. Some accessions were ranked high or low for WUE at one temperature stage only, whereas others were ranked high or low for WUE across temperatures. For example, PI550604 was among the bottom-ranked accessions for WUE at each temperature, and PI534165 was among the bottom-ranked at 24°C and 32°C. PI455221 was a top-ranked accession for WUE at both 15°C and 24°C, while PI329403 was among the highest-ranked accessions at 15°C and 32°C.

**Figure 5.**
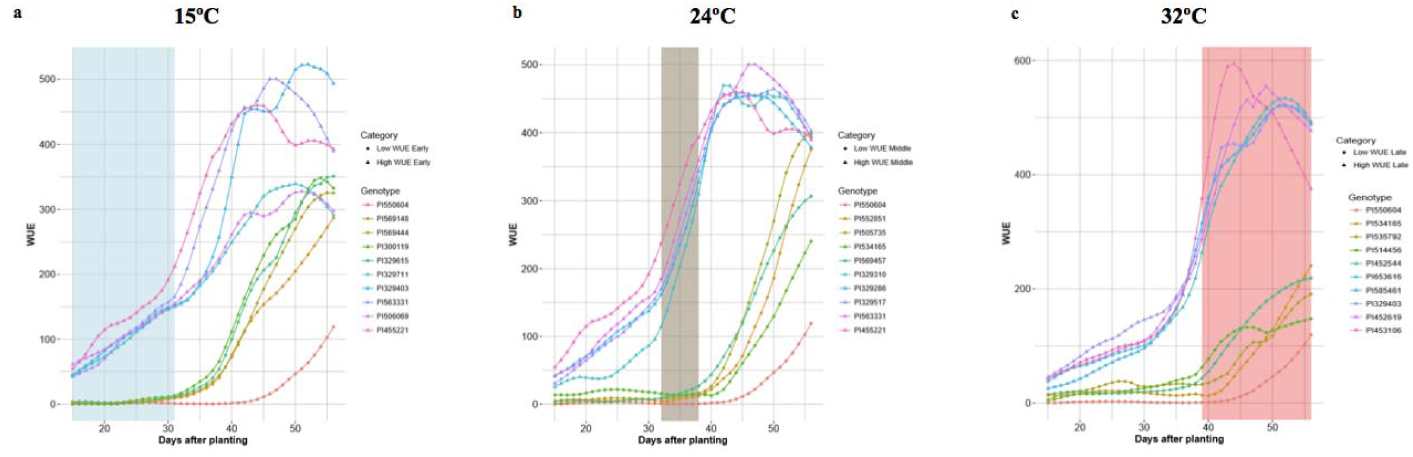
Temporal Profiles of WUE for the Highest and Lowest WUE Accessions at Each Temperature Phase. Each panel depicts the five highest and five lowest WUE accessions at each temperature phase. The colored lines show the changes in WUE for each of those accessions throughout the experiment as the plant developed and temperature changed. (a) WUE profiles for the five highest and the five lowest WUE accessions at 15°C. (b) WUE profiles for the five highest and the five lowest WUE accessions at 24°C. (c) WUE profiles for the five highest and the five lowest WUE accessions at 32°C.

For each accession, we analyzed the profile of each phenotype over time and at different temperatures. Again, for brevity, here we only refer to WUE. For example, the temporal WUE profiles for the BAP accessions separate into seven clusters, each represented by a growth curve represented by the average of all accessions in that cluster (Fig. 6a). The temporal WUE profile for each of the seven groups is distinct. The mean WUE for accessions that comprise Cluster 1 is low at 15°C, shows a minimal increase at 24°C, and increases steadily at 32°C, but never reaches levels as high as several other clusters. Cluster 2 shows a slightly higher WUE at 15°C compared to Cluster 1 and an increase in WUE at 24°C and 32°C, but never exceeds moderate levels of WUE. In Cluster 3 there is little increase in WUE until the temperature increases to 32°C and this cluster eventually reaches moderate WUE. WUE for Cluster 4 is low at 15°C, increases at 24°C and 32°C. Cluster 5 exhibits a modest increase in WUE at 15°C, a rapid increase at 24°C, but WUE increases slowly at 32°C. Cluster 6 has an early and robust increase in WUE across all three temperatures in the experiment and reaches the maximum peak WUE of all seven clusters. Finally, WUE for Cluster 7 is moderate at 15°C and increases steadily at 24°C and 32°C, but ultimately peaks at a moderate level. Cluster analysis for each additional trait is in S7. The dendrogram tree (Fig. 6b) depicts the hierarchical clustering of the BAP accessions into seven groups based on their temporal WUE profiles, with each cluster represented by a different color. The clustergram of the PCA-weighted mean (Fig. 6c) illustrates the divergence of the temporal WUE profiles of the BAP accessions into the seven stable clusters which captured the maximum phenotypic difference for this trait.

**Figure 6.**
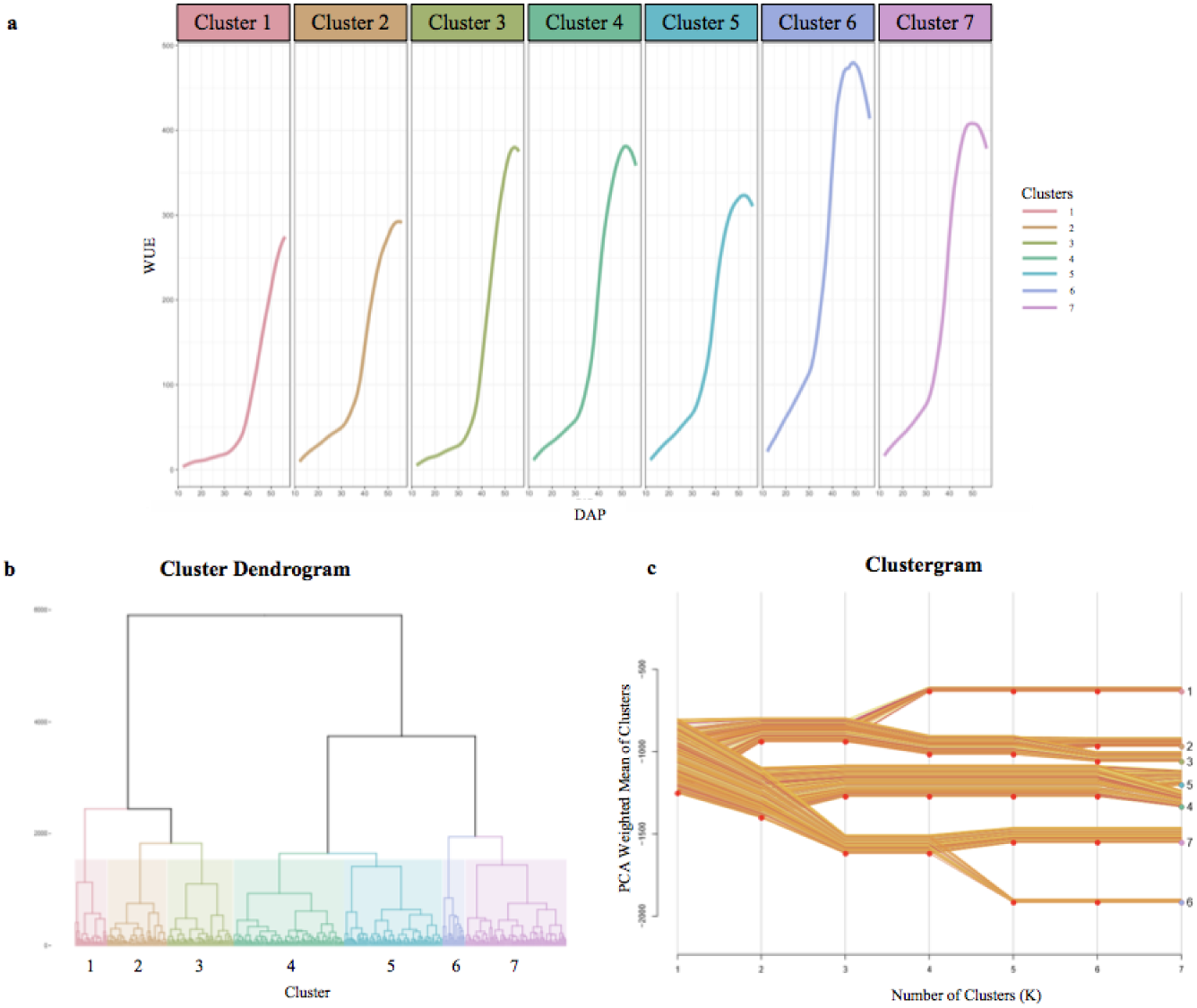
BAP Accessions Separate into Seven Distinct Temporal WUE Clusters. (a) The temporal WUE profiles of BAP accessions from 12 DAP to 56 DAP separate into seven clusters. The growth curves represent the average WUE profile over time for each cluster. (b) Hierarchical dendrogram tree showing the segregation of the BAP accessions into 7 clusters, each represented with a different color. (c) Clustergram of the PCA-weighted mean of each cluster showing the divergence of the BAP accessions into the seven clusters, and the distances between the clusters. The cluster numbers and colors correspond to the dendrogram (b) and temporal WUE profiles (a).

### Genome-Wide Association Study

A genome-wide association study (GWAS) was performed to characterize the genetic architecture underlying response to early cold stress conditions in the BAP. GWAS was performed using the multi-locus mixed model (MLMM) algorithm (11). MLMM uses multiple loci in the model to yield a higher detection power and lower the potential of false discoveries. Figure 7 illustrates the detection of highly significant signals related to each trait of interest on all ten sorghum chromosomes. Each significant SNP has a maximum corrected p-value of 0.05. This analysis identified 2,305 highly significant SNPs associated with biomass, height, hull area, WUE, and RGR.

**Figure 7.**
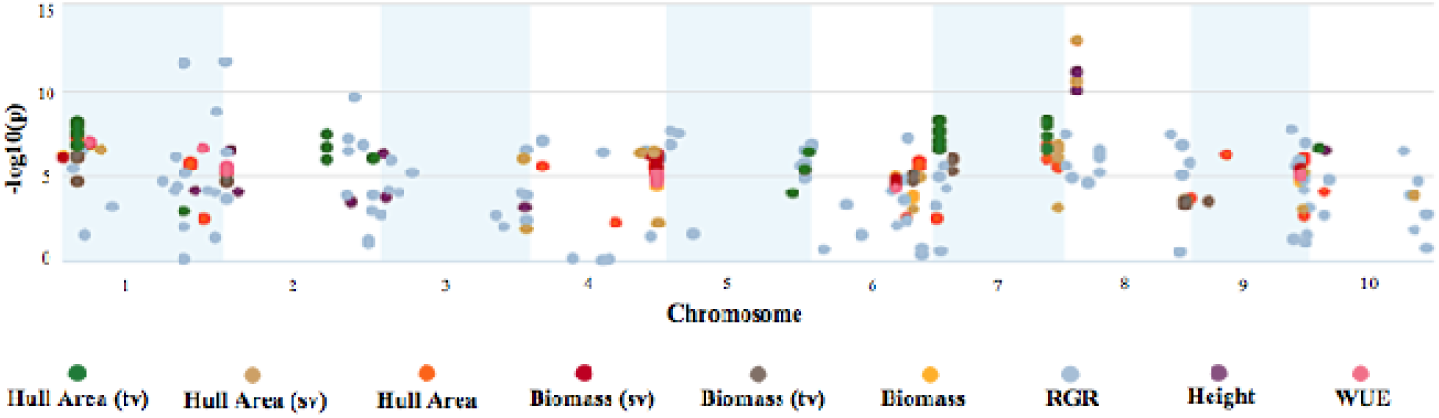
Manhattan Plot of GWAS on Biomass, Growth, and WUE Traits. The Manhattan plot depicts significant SNPs for each trait of interest and the corresponding -log_10_(p) of the GWAS mixed linear model with no other SNPs included in the model.

To identify SNPs significantly correlated with phenotypes at a specific DAP and correlated with temperature changes, we also used the values for each phenotype at each DAP (Fig. 8) for the GWAS.

**Figure 8.**
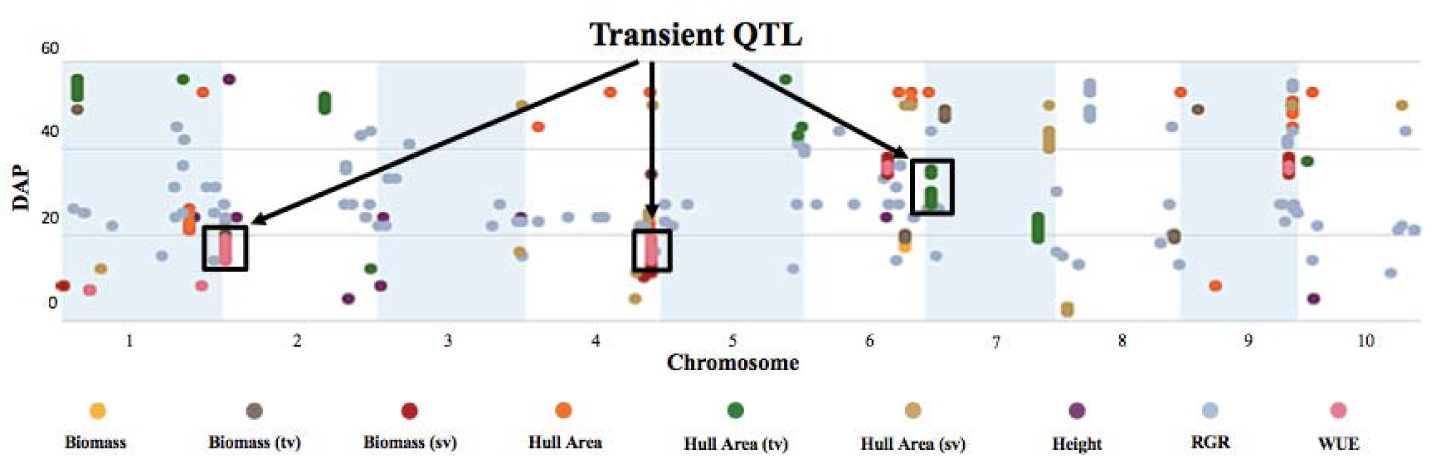
Temporal Manhattan Plot of GWAS on Biomass, Growth, and WUE Traits Reveals Transient QTL. The plot depicts highly significant SNPs corresponding to the traits of interest at specific DAP. The y-axis denotes the DAP, and the x-axis represents the sorghum chromosome. Black boxes highlight transient QTL that “turn on/off” with developmental stage and as the temperature increases.

This analysis revealed several QTL that displayed a transient response, appearing at specific times and developmental stages, and as temperatures increased. (Fig. 8). For example, the highly significant hull area SNP 7:2,934,702 was transiently detected from 27 to 30 DAP and again from 34 to 35 DAP (Fig. 9).

**Figure 9.**
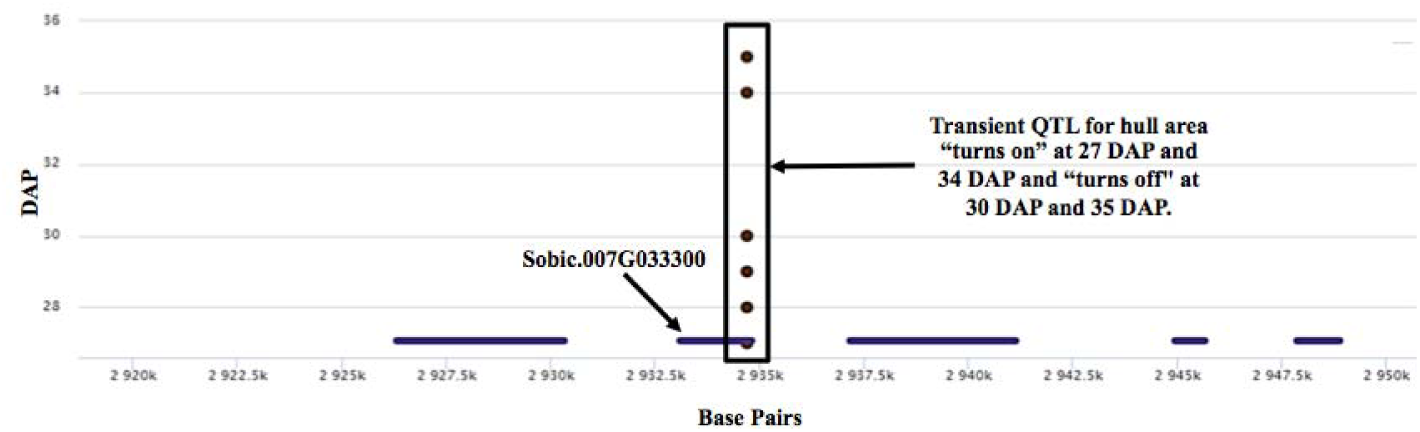
A Transient QTL Associated with the Hull Area Trait. The boxed region represents a transient hull area QTL at SNP 7:2,934,702 that “turns on/off” with developmental stage and as temperature increases. There are five genes within 15kb of the transient QTL. SNP 7:2,934,702 is directly on candidate gene Sobic.007G033300.

For the transient QTL that were detected, we also examined the phenotypic differences for the alternative alleles at a particular SNP. For the hull area SNP 7:2,934,702 shown in Figure 9, allele boxplots (Fig. 10, S8) shows that a “G” at the position corresponds to a significantly higher value for hull area than a “C” for 27 to 30 DAP and 34 to 35 DAP. SNP 7:2,934,702, within gene Sobic.007G033300, encodes plastocyanin-like protein. Plastocyanins are copper-containing chloroplast-localized proteins involved in photosynthesis and function in electron transfer between photosystem II and photosystem I (12). This putative plastocyanin, Sobic.007G033300, is highly expressed in sorghum shoots, leaves, and early inflorescences (https://phytozome.jgi.doe.gov/pz/portal.html) (13). We examined other sequence variation existing within this gene using public re-sequencing data and found seven SNP variants resulting in nonsynonymous (NS) amino acid substitutions and an additional inframe three basepair deletion segregating within individuals in the BAP (S9a). Eighty of the phenotyped BAP accessions contain no variants in Sobic.007G033300 relative to the reference sequence, while the remaining accessions contain between one to eight of the variants summarized in S9a. When grouped, individuals carrying at least one variant in Sobic.007G033300 have a significantly larger hull area from 27 DAP compared to accessions containing no variants (S9b).

**Figure 10.**
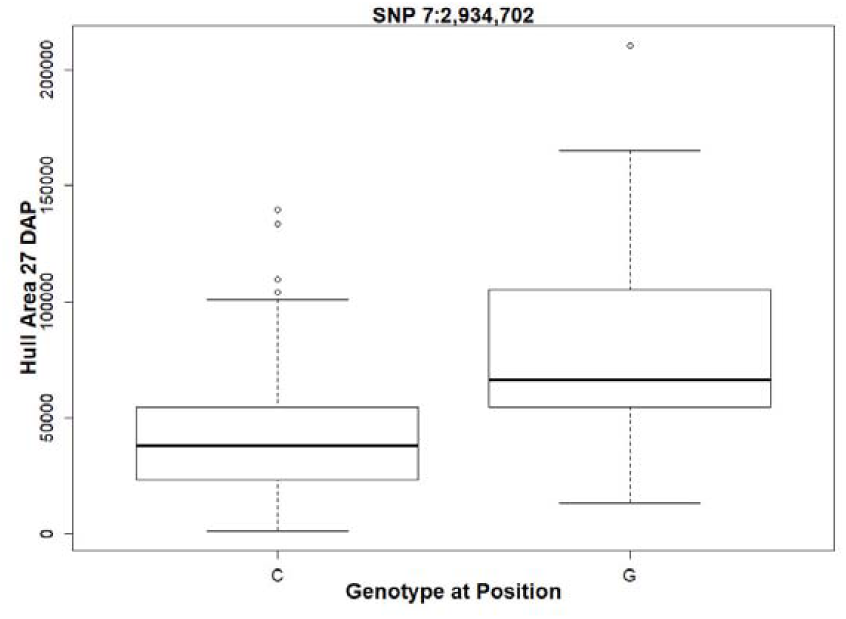
Phenotypic Differences in Hull Area Associated with a SNP in a Plastocyanin-like Gene. The allele boxplot shows the correlation of a “G” at SNP 7:2,934,702 with a significantly larger hull area at 27 DAP. S8 contains additional boxplots showing this correlation for 28-30 DAP and 34-35 DAP.

The GWAS also identified pleiotropic QTL. A transient QTL on chromosome two (SNP 2:1,199,928) was significant for WUE and biomass from 14 to 19 DAP. SNP 4:64256235 on chromosome four was a significant transient QTL for WUE and biomass from 14 to 19 DAP, and height at 34 DAP. Transient QTLs on chromosomes six and nine (SNP 6:42,116,590 and SNP 9:55067080, respectively) were significant for WUE and biomass at 35 to 36 DAP. S10 shows the traits and DAP associated with each SNP.

### Candidate Gene Identification

We conducted a genome scan of the 15 kb upstream and 15 kb downstream of each significant SNP to identify candidate genes. Functional annotations, putative homologs, and polymorphisms within the candidate genes based on public genomic re-sequencing data were analyzed using Phytozyme (https://phytozome.jgi.doe.gov/pz/portal.html). GWAS identified highly significant QTL near 72 candidate genes (S11) with putative functions potentially related to biomass, height, hull area, WUE, and RGR, and in response to cold. Many of these candidate genes are close to pleiotropic QTL implicated in cold stress response in addition to the biomass-associated phenotypes.

For brevity, here we will only refer to one *a priori* candidate gene. GWAS identified SNP 6:40,312,463 which was significantly associated with RGR at 33 DAP and is located within the gene Sobic.006G057866, which encodes *PSEUDO-RESPONSE REGULATOR 7 (PRR37) / MATURITY 1 (Ma1)* and is involved in flowering time regulation in sorghum (14). SNP 6:40,312,463 causes a non-synonymous amino acid substitution (Asn184Lys) within *Ma1*, and although accessions carrying this polymorphism show a slightly reduced mean RGR at 33 DAP, they were not significantly different (S12a). This observation suggests that allelic variation at SNP 6:40,312,463 is not likely to be a robust contributor to phenotypic variation in RGR. However, interestingly, most accessions (∼90%) including both BAP accessions and lines from other diversity panels carry a polymorphism in *Ma1* resulting in loss of the annotated reference stop codon, which could drastically increase the protein size and affect its function (S12b).

## DISCUSSION

Sorghum is attractive as a bioenergy feedstock crop because it is heat and drought tolerant and can thrive in marginal environments. It is an ideal target for accelerated improvement through breeding and engineering because it has extensive genetic and phenotypic diversity but has not yet benefited from genomics or genetic modification like some other crops such as maize. Early season planting of sorghum provides the opportunity for an extended growing season with higher potential accumulation of biomass. However, as a tropical crop, sorghum is sensitive to cold stress. Identification of accessions that exhibit the highest WUE, RGR, height, hull area, and biomass under early cold stress conditions could facilitate genetic improvement of bioenergy sorghum for early season planting and cultivation in colder temperatures.

Our results identified the top performing BAP accessions for bioenergy-related traits under early cold stress. We identified the accessions with the highest and lowest rankings for each trait and multiple bioenergy-related traits. Accession PI329299 was among the top accessions for hull area and height, PI452619 was a top accession for hull area and WUE, PI329403 was a top accession for hull area and WUE, and PI585461 was among the top accessions for both biomass and WUE. These top-performing accessions, particularly the accessions that possess multiple advantageous traits, are promising candidates for sorghum bioenergy breeding programs and the development of additional genetic resources such as mapping populations (e.g., NAMs).

We defined ideotype-positive and ideotype-negative accessions as accessions that ranked high or low, respectively, for multiple traits or exhibited beneficial trait combinations. The ideotype-positive phenotype is the combination of high WUE with a low height-to-biomass ratio, i.e., a plant that achieves high biomass but does not grow tall. This ideotype is desirable for bioenergy sorghum because the required biomass accumulation is attained with reduced water use and without the increased risk of lodging associated with taller plants. Five accessions were ideotype-positive in the early phase of the experiment when the temperature was 15°C. These accessions may be beneficial for breeding bioenergy sorghum that can be planted early in the season or grown in colder environments. Accession PI329517, which attained the ideotype-positive phenotype early under cold stress, and maintained it after the temperature increased, may be beneficial for early season planting, for long growth periods, or under conditions of reduced water availability. In contrast, ideotype-negative accessions that have relatively low biomass accumulation, ranked low for WUE, and are tall may be undesirable for bioenergy sorghum because more water would be required and with a higher risk of lodging and decreased biomass accumulation. Even though these accessions may not be desirable for breeding, identification of these extreme ideotypes may be useful for the development of structured populations for further genetic analysis.

The profiles of the traits over time and in response to temperature changes could be discerned, ranked, and clustered using the phenotypic data collected with single-day resolution. The BAP clustered into 6 to 8 distinct temporal profiles for each trait, identifying accessions that perform best under early cold stress, and at different temperatures and development stages. For example, accessions in WUE clusters two, six, and seven have the highest WUE at 15°C. Similarly, cluster six not only attains high WUE under 15°C but is the cluster with the highest WUE attained under increased temperatures and therefore may perform well under drought conditions. Accession PI455221 was among the top-performing accessions for WUE at both 15°C and 24°C, and PI329403 was among the top-performing accessions for WUE at 15°C and 32°C, demonstrating that high WUE can be maintained as the plants develop and as the temperature increases.

GWAS revealed 2,305 highly significant SNPs associated with biomass, hull area, WUE, RGR, and height phenotypes. We also identified significant SNPs that colocalized for multiple traits, suggesting that these polymorphisms may be near tightly linked genes or genes with pleiotropic effects. By using the daily values for each phenotype in the GWAS, we determined the temporal correlations of SNPs to traits. With this approach, we identified transient QTL that “turn on/off” at a specific developmental stage or with a change in temperature.

The highly significant SNPs identified by GWAS revealed 72 candidate genes with putative functions related to the bioenergy-relevant and cold-stress-responsive traits of interest. A transient QTL for RGR at 33 DAP mapped to the gene Sobic.006G057866, which encodes *PRR37/Ma1* an *a priori* candidate gene that represses flowering in long days (14). The observation that this transient QTL for RGR “turns on” at 33 DAP, a critical time for the transition to flowering in sorghum, and immediately after the temperature increased from 15°C, is consistent with a role for *PRR37/Ma1* in growth and biomass accumulation through regulation of the transition from vegetative to reproductive development. Another transient QTL associated with hull area was significant from 27 to 30 DAP and again from 34 to 35 DAP. This transient QTL mapped to the gene Sobic.007G033300, which encodes a putative plastocyanin that may function in photosynthetic electron transfer. Additional sequence variation in this gene correlated with a significantly larger hull area from 27 to 30 DAP. These observations suggest that Sobic.007G033300 is important for biomass accumulation, consistent with known deleterious effects of cold on photosynthesis, in particular, the capacity for electron transport and photosystem protection (15). Studies have also implicated that plastocyanins affect yield in crops such as rice (16).

In summary, we tracked growth over time and under early season cold temperature stress in a genetically diverse sorghum population using daily image-based phenotyping. We analyzed the changes in traits over time, which would not have been possible with the collection of endpoint phenotypes alone. The BAP segregated into distinct clusters for each trait, identifying accessions that performed best under early cold stress, and at different temperatures and development stages. GWAS on the daily phenotypes revealed transient QTLs for highly heritable bioenergy-relevant traits. In some cases, we could identify putative causative alleles in candidate genes that correlated to a significant difference in phenotype. These findings may facilitate targeted genetic modifications or genomics-driven breeding efforts for the improvement of bioenergy sorghum.

## METHODS AND MATERIALS

### Plant Materials

This study used the sorghum Bioenergy Association Panel (BAP) (1). Details about the panel design, GBS genotyping, marker distribution, population structure, and linkage disequilibrium (LD) decay have been previously described (1). We selected 369 BAP accessions genotyped at 232,303 SNPs. The BAP accessions represent a racially, geographic, and phenotypically diverse selection of sorghum accessions, but is limited to accessions exhibiting key bioenergy traits, such as height, sensitivity to photoperiod, and delayed flowering.

### Experimental Conditions

Three replicates of 369 BAP accessions (1131 plants) were planted in 600 g of Turface® in 8-inch tall tree pots, with +14-14-14 Osmocote (1.5 lb/cubic yard) fertilizer. The potted seeds were held overnight in a growth chamber at 32°C (day)/22°C (night). The next day, the pots were loaded into a carrier on an automated phenotyping system within a controlled-environment plant growth chamber. The phenotyping system moved the plants on a closed-loop conveyor path to stations for daily watering, weighing, and imaging. The plants were positioned in the growth chamber in a randomized block design and were rotated one-half lane each day to reduce edge effects. To study the impact of cold stress on these accessions, plants were grown at 15°C (day)/15°C (night) for 31 days, 24°C (day)/19°C (night) for 7 days, and 32°C (day)/22°C (night) for 18 days. The soil was maintained at 100% field water capacity by watering the plants twice daily to a target mass of 1192 grams. The target mass was calculated by adding the mass of the plant carrier (342g), the water mass at saturation (250g), and the mass of the Turface®-filled pot (600g). Water was added after each potted plant in its carrier was weighed, and the volume of water added to reach the target mass of 1192 grams was recorded.

### Image Collection and Phenotyping

Each plant passed through a visible light imaging chamber daily while on the automated phenotyping system. The imaging cameras recorded two side views (sv) and one top view (tv) image. As the plants grew, the fields of view on the cameras were adjusted so that the entire plant could be captured in each image. The optical zoom level was reduced for the top view and side view images at 19 DAP and 40 DAP. Scaling factors were calculated for both area and height using a reference object of known size, so that pixel areas across zoom levels were comparable. After eight weeks (56 DAP), the plants were removed from the phenotyping system. The shoot of each plant was cut at the base of the stem. Fresh weight measurements of the shoots were collected immediately.

The images were analyzed using Plant Computer Vision (PlantCV), an image-processing tool coded in Python (17). The pixel areas from the daily top view and two side view images of each plant were analyzed with PlantCV to generate measurements of the area, hull area, height, RGR, and WUE.

The area was calculated by adding the pixel count from the top view image to the two side view images. Endpoint fresh weight measurements were correlated with area calculations at 51 DAP to estimate biomass (17) (S13). The values at 51 DAP were used for the correlation because, after that date, plants began to overlap and grow outside of the camera’s field of view, and the resulting pixel counts were less indicative of the actual plant size.

The hull area is the convex hull calculated from the pixel count in the smallest area that includes a set of given points in a plane. WUE was calculated by dividing the derived area by the cumulative water added to each plant. Height was determined from the side view images.

The relative growth rate (RGR) was calculated as described in Hoffmann and Poorter, using estimator 2 to determine the RGR for each distinct time point (18). Briefly, we calculated the natural log of all replicate area values in the experiment, then calculated the mean of the log-transformed values for two time points, t_1_ and t_2_. The mean value of the log-transformed areas for t_2_, W2, was subtracted from the mean value of the log-transformed areas for t_1_, W_1_, and then divided by the difference between t_2_ and t_1_ as shown:

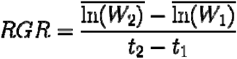

Germination dates were determined for each plant based on the top view area measurements. The earliest DAP with a top view area measure was considered the germination date. The resulting germination date for the three replicates of each accession was averaged to determine the germination date for each accession.

### Data Processing and Analysis

Image-derived phenotypic data were generated for 309 of the BAP accessions (S1). Phenotypic analysis was performed on the image-derived data using the R statistical software (R). Plants that never germinated or that died by 46 DAP were excluded from the analysis. A conservative initial outlier removal step was performed on each phenotype to remove likely artifacts from the image analysis. A data point was considered an outlier and excluded from further analysis if its value was greater than 40 median absolute deviations (MAD) from the median (19). Raw measurements from PlantCV were smoothed using predicted values from a loess smoothing fit. These fitted values were used as the phenotypes for further analyses (20).

The ‘lmer’ function in the lme4 R package was used to estimate variance components for broad-sense heritability (21). Heritability was calculated based on the subset of accessions that had three replicates that germinated. Broad-sense heritability was calculated as:

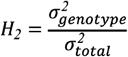

After the heritability calculation, the median value of the replicates for each accession was used for further analysis. Traits were tested for normality and transformed as necessary using the Box-Cox procedure as implemented in R with the ‘boxcox’ function in the MASS package (22).

### Hierarchical Clustering

Hierarchical clustering of each trait from 12 to 56 DAP was performed using the ‘hclust’ R function using a Euclidean distance matrix and the Ward agglomeration method (23). Clustering results were visualized both as a dendrogram using the R package ‘dendextend’ and as a clustergram using the function ‘clustergram.R’ (24, 25).

### Ideotype Selection

Height-to-biomass ratios were calculated for each day and converted to a percentile with values between 0 and 100. The average percentile of the accessions for height-to-biomass ratio and WUE over the growing period, DAP 15 to 31 or DAP 39 to 56, for the early and late periods, respectively, was used to select ideotype accessions. Ideotype-positive accessions were defined as those with a height-to-biomass ratio in the bottom 5 percent of the population and a WUE in the top 10 percent of the population. Ideotype-negative accessions have a height-to-biomass ratio in the top 5 percent of the population and a WUE in the bottom 10 percent. Radar plots of the ideotype accessions were generated with the R package ‘fmsb’ (26). Heatmaps were generated using the R package ‘pheatmap’ (27).

### Genome-wide Association Study

Genotyping-by-sequencing (GBS) SNP markers for the BAP have been previously described (1). GWAS was performed using a multi-locus mixed linear model (MLMM) in R to identify loci associated with each trait of interest. The first three principal components of the genotype matrix were included as covariates in the mixed model to control for population structure (28). A kinship matrix, calculated from the genotype matrix using the Astle-Balding method in the ‘synbreed’ package, was also included as a random effect to control for familial and cryptic relatedness between accessions (29, 30) (S2).

MLMM tests for association with the phenotype using a stepwise mixed model regression. In each step, the SNP with the most significant association to the phenotype is added to the model as a covariate. Stepwise addition of SNPs continues until the heritable variance estimate (pseudo-heritability) reaches 0 (11). A final set of high-confidence SNPs were selected as those that were included as covariates in either of the two optimal models chosen by the MLMM software: the multiple-Bonferonni model or the extended BIC model (11). In the multiple-Bonferroni model, all cofactors with a p-value below a Bonferroni corrected threshold are selected. Multiple-Bonferroni was the more stringent of the two models (11). The extended BIC model selects a model based upon BIC penalized by the model complexity (1, 31).

The temporal GWAS results were analyzed using ZBrowse, an interactive browser that runs using R (32). Using ZBrowse, we were able to view the SNPs for multiple traits simultaneously and plot those traits over time to determine the DAP on which each SNP was significant and thus identify transient QTL peaks that turn “on/off” at specific DAP.

### Candidate Gene Identification

The BTx623 sorghum reference genome version 3.1 was used to identify genes colocalizing with or adjacent to the associated SNPs. A genome scan of 15 kb upstream and 15 kb downstream of each significant SNP was performed to identify candidate genes. Phytozome (https://phytozome.jgi.doe.gov/pz/portal.html) was used to analyze the functional annotation of the candidate genes and to identify putative homologs in other species. Analysis of polymorphisms within candidate genes was done using three diversity panels of re-sequencing data available for sorghum via Phytozome (33-35). VCFtools was used to generate final variant calls after merging the three diversity VCFs, based on a minor allele frequency cutoff of >0.05, and to create distinct VCFs for the accessions used in this study (36). Variant effects were estimated using the SNPeff pipeline and the v3.1 sorghum reference to assess the potential impact of sequence variants on annotated gene models (37).

## Supporting information

Supplemental References

## ACKNOWLEDGMENTS

For their assistance with data collection, we thank John Gierer, Skyler Mitchell, Darren O’Brien, Phil Ozersky, Brandon Patrick, Dr. Miranda Haus, and Madeline Wiechert. We also thank Mindy Darnell and Leonardo Chavez from the Donald Danforth Plant Science Center’s Bellwether Phenotyping Facility. This research was funded in part by the Advanced Research Projects Agency-Energy (ARPA-E), U.S. Department of Energy, under Cooperative Agreement Number DE-AR0000594. The views and opinions of authors expressed herein do not necessarily state or reflect those of the United States Government or any agency thereof.

## AUTHOR CONTRIBUTIONS

NS and TM conceived and designed the project. NS guided the process of analyses. NF and CL derived the phenotypic values from the image data. GZ performed GWAS. GZ and SL performed computational analyses. SL performed genomic analyses. EA, GZ, SL, and NS performed phenotypic data analyses and interpretation. EA, GZ, SL, IB, NS, and TM wrote the paper. All authors read and approved the manuscript.

## SUPPLEMENTARY FILES

**S1**. Table of Bioenergy Association Panel accessions used in this study (adapted from Brenton et al., 2016) with image-derived phenotypic data.

**S2**. Heatmap of a kinship matrix showing correlation analysis among the 369 BAP accessions. The color histogram shows the distribution of coefficients of coancestry values in the whole kinship matrix. The color scale shows the degree of correlation (white-yellow, low correlation; orange-red, strong correlation). C1: 25/29 accessions from Ethiopia, 26/29 photoperiod-sensitive. C2: 47/48 accessions from Ethiopia. C3: All photoperiod-insensitive accessions. C4: All cellulosic accessions; 29/30 photoperiod-sensitive.

**S3**. Table of Germination Data. (a) Accessions with an average germination time within 10 DAP. (b) Plants that did not germinate. (c) Average germination times of accessions that germinated after 10 DAP, calculated from the three replicates for each BAP accession.

**S4**. Heritability of Traits.

**S5**: Table ranking accessions for each trait at 51 DAP.

**S6**. Table ranking accessions for each trait at each temperature.

**S7**: Cluster analysis for biomass, height, hull area, and RGR.

**S8**: Allele Boxplots show the correlation of a “G” at SNP 7:2,934,702 with a significantly larger hull area for 27-30 and 34-35 DAP.

**S9**. (a) The table shows all variants resulting in non-synonymous amino acid substitutions within the coding sequence of Sobic.007G033300 within BAP accessions and summarizes the potential impacts of variants based on SNPeff predictions. Variants with minor allele frequency < 5% were removed. ^**i**^ Denotes the percentage of accessions within the BAP that carry the particular variant. (b) Boxplot showing hull area (27 DAP) for BAP accessions based on the presence of 1 to 8 moderate effect SNPs in gene Sobic.007G033300 resulting in non-synonymous amino acid substitutions (Alternate) versus accessions with no variants compared to reference (Reference). The analysis excluded accessions containing heterozygous alleles. Alternate accessions have significantly larger hull area (p = 0.019) compared to reference accessions according to Welch’s *t*-test.

**S10**: Table of SNPs with correlated traits.

**S11**: Table of 72 candidate genes.

**S12**. (a) Boxplot showing the relative growth rate for individuals within the BAP based on the presence of a missense allele resulting in a non-synonymous amino acid substitution (Chr 6:40312463, Asn184Lys). Accessions heterozygous at the SNP were not different according to Welch’s t-test.

The table shows all variants within the coding sequence of Sobic.006G057866 (PRR37/Ma1) in the BAP accessions and summarizes the potential impacts of variants based on SNPeff predictions. Variants with minor allele frequency < 5% were removed. ^**I**^ Denotes the percentage of accessions within the BAP that carry the particular variant.

**S13**. Scatterplot and regression show the correlation of area at 51 DAP to endpoint fresh weight.

