## Supplementary figures and images for "Genomewide association study reveals transient loci underlying the genetic architecture of biomass accumulation under cold stress in Sorghum"

### Supplemental File 2

#### Slide 1
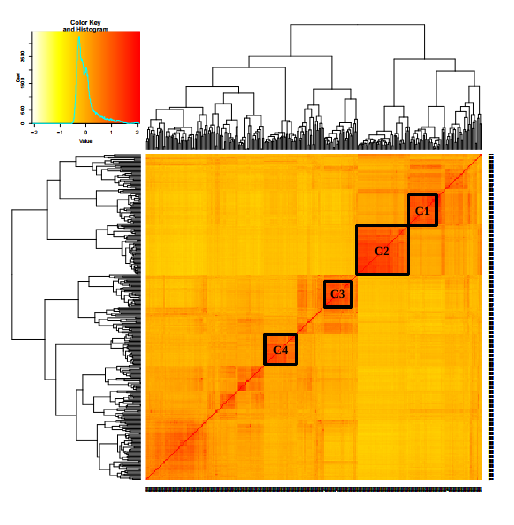

C1
C2
C3
C4

### Supplemental File 8

#### Slide 1
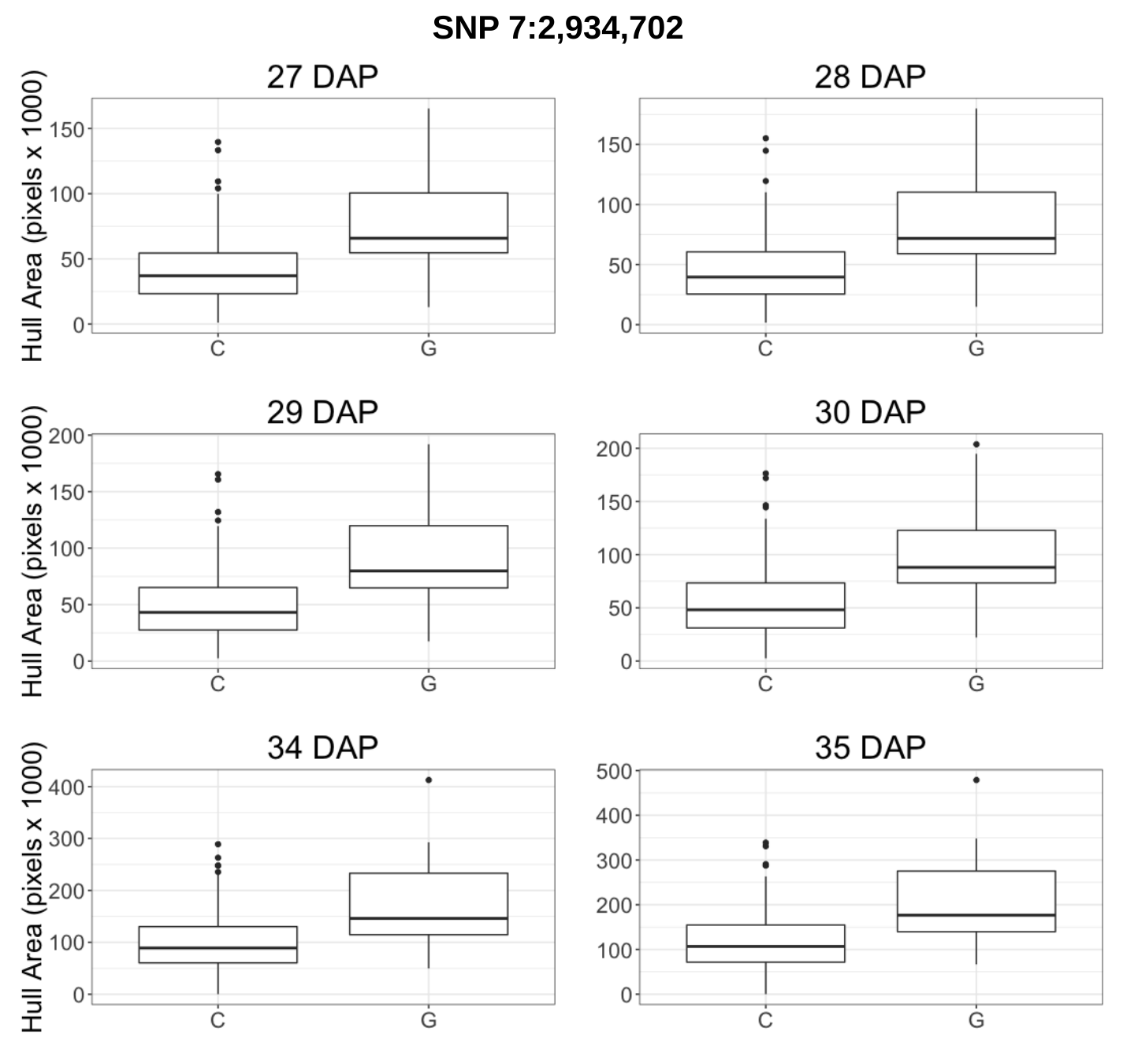

SNP 7:2,934,702

### Supplemental File 13

#### Slide 1
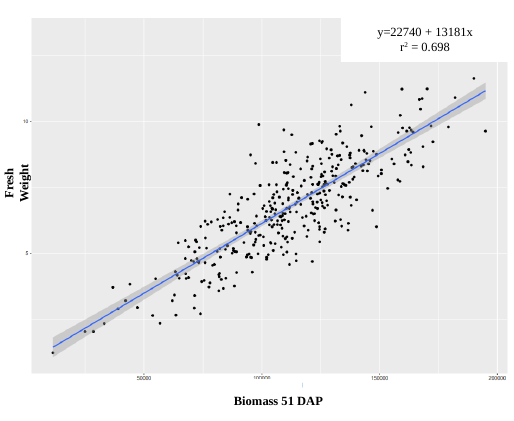

y=22740 + 13181x
r^2^ = 0.698
Fresh Weight
Biomass 51 DAP
