## Supplemental File 7 for "Genomewide association study reveals transient loci underlying the genetic architecture of biomass accumulation under cold stress in Sorghum"

### Slide 1
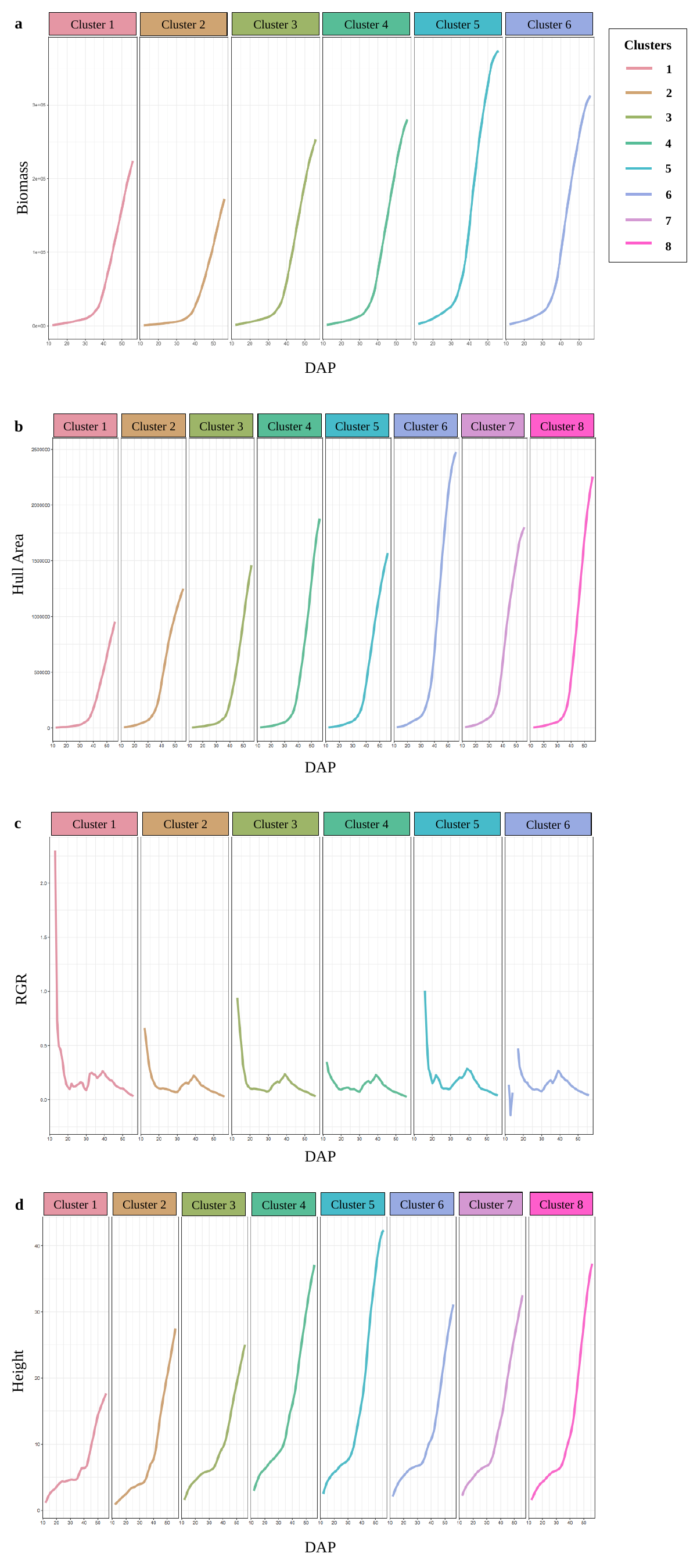

a
Cluster 6
Cluster 5
Cluster 4
Cluster 3
Cluster 1
Cluster 2
Clusters
1
2
3
4
5
Biomass
6
7
8
DAP
b
Cluster 1
Cluster 7
Cluster 8
Cluster 2
Cluster 3
Cluster 4
Cluster 5
Cluster 6
Hull Area
DAP
c
Cluster 5
Cluster 3
Cluster 4
Cluster 1
Cluster 2
Cluster 6
RGR
DAP
d
Cluster 8
Cluster 7
Cluster 1
Cluster 2
Cluster 5
Cluster 6
Cluster 3
Cluster 4
Height
DAP
