## Supplemental File 9 for "Genomewide association study reveals transient loci underlying the genetic architecture of biomass accumulation under cold stress in Sorghum"

### Slide 1
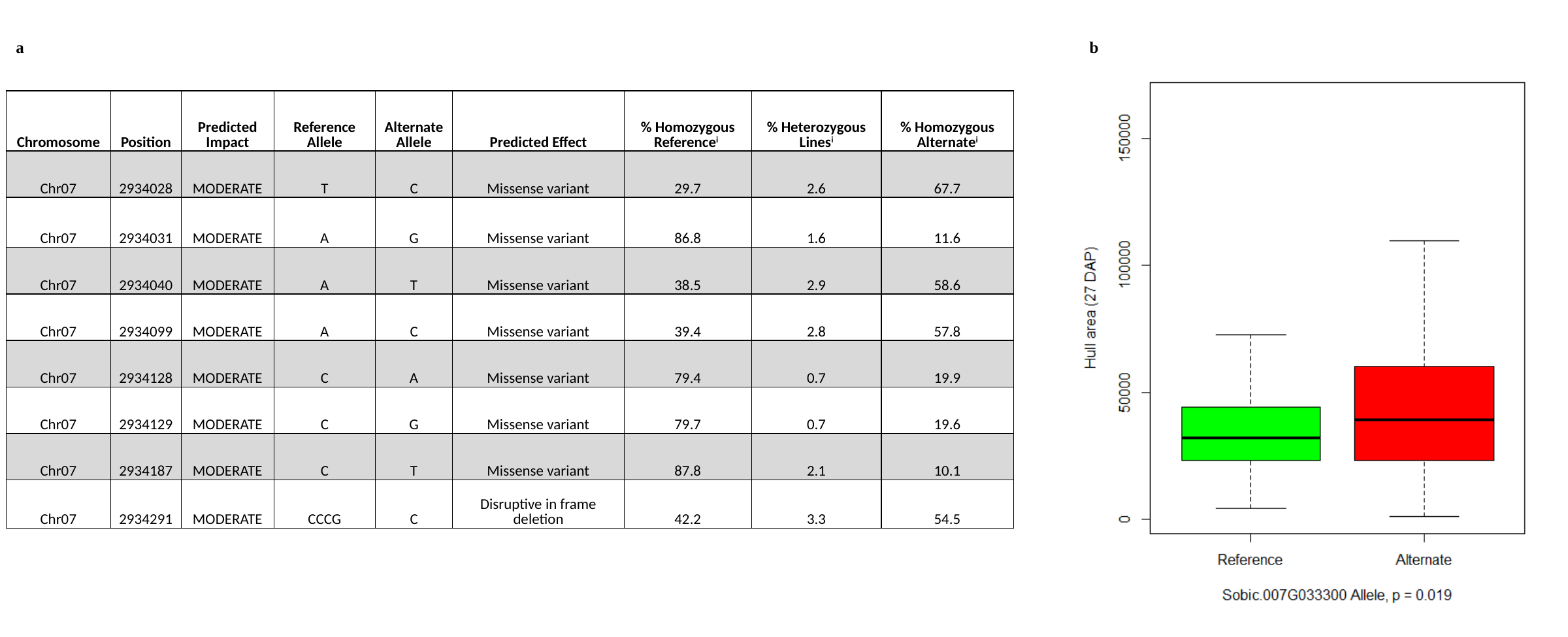

a
b
| Chromosome | Position | Predicted Impact | Reference Allele | Alternate Allele | Predicted Effect | % Homozygous Reference^i^ | % Heterozygous Lines^i^ | % Homozygous Alternate^i^ |
| --- | --- | --- | --- | --- | --- | --- | --- | --- |
| Chr07 | 2934028 | MODERATE | T | C | Missense variant | 29.7 | 2.6 | 67.7 |
| Chr07 | 2934031 | MODERATE | A | G | Missense variant | 86.8 | 1.6 | 11.6 |
| Chr07 | 2934040 | MODERATE | A | T | Missense variant | 38.5 | 2.9 | 58.6 |
| Chr07 | 2934099 | MODERATE | A | C | Missense variant | 39.4 | 2.8 | 57.8 |
| Chr07 | 2934128 | MODERATE | C | A | Missense variant | 79.4 | 0.7 | 19.9 |
| Chr07 | 2934129 | MODERATE | C | G | Missense variant | 79.7 | 0.7 | 19.6 |
| Chr07 | 2934187 | MODERATE | C | T | Missense variant | 87.8 | 2.1 | 10.1 |
| Chr07 | 2934291 | MODERATE | CCCG | C | Disruptive in frame deletion | 42.2 | 3.3 | 54.5 |
