## Supplemental File 12 for "Genomewide association study reveals transient loci underlying the genetic architecture of biomass accumulation under cold stress in Sorghum"

### Slide 1
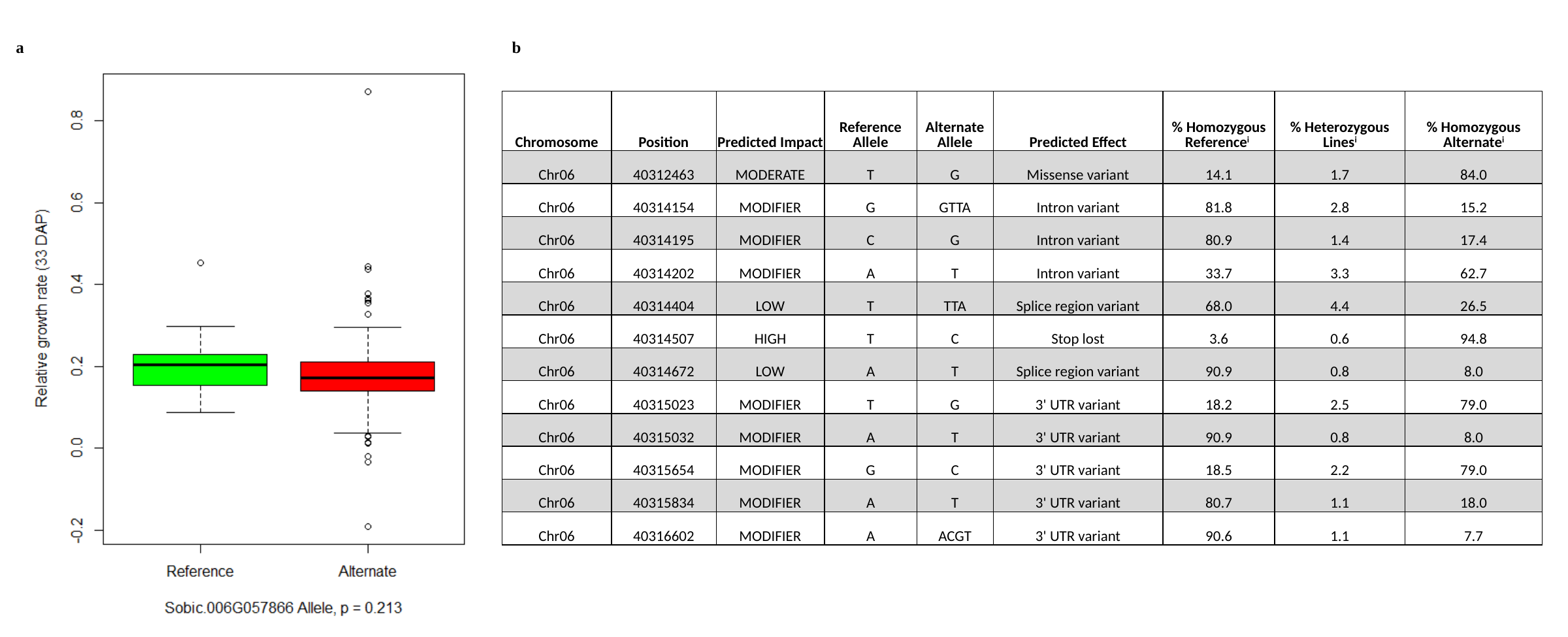

a
b
| Chromosome | Position | Predicted Impact | Reference Allele | Alternate Allele | Predicted Effect | % Homozygous Reference^i^ | % Heterozygous Lines^i^ | % Homozygous Alternate^i^ |
| --- | --- | --- | --- | --- | --- | --- | --- | --- |
| Chr06 | 40312463 | MODERATE | T | G | Missense variant | 14.1 | 1.7 | 84.0 |
| Chr06 | 40314154 | MODIFIER | G | GTTA | Intron variant | 81.8 | 2.8 | 15.2 |
| Chr06 | 40314195 | MODIFIER | C | G | Intron variant | 80.9 | 1.4 | 17.4 |
| Chr06 | 40314202 | MODIFIER | A | T | Intron variant | 33.7 | 3.3 | 62.7 |
| Chr06 | 40314404 | LOW | T | TTA | Splice region variant | 68.0 | 4.4 | 26.5 |
| Chr06 | 40314507 | HIGH | T | C | Stop lost | 3.6 | 0.6 | 94.8 |
| Chr06 | 40314672 | LOW | A | T | Splice region variant | 90.9 | 0.8 | 8.0 |
| Chr06 | 40315023 | MODIFIER | T | G | 3' UTR variant | 18.2 | 2.5 | 79.0 |
| Chr06 | 40315032 | MODIFIER | A | T | 3' UTR variant | 90.9 | 0.8 | 8.0 |
| Chr06 | 40315654 | MODIFIER | G | C | 3' UTR variant | 18.5 | 2.2 | 79.0 |
| Chr06 | 40315834 | MODIFIER | A | T | 3' UTR variant | 80.7 | 1.1 | 18.0 |
| Chr06 | 40316602 | MODIFIER | A | ACGT | 3' UTR variant | 90.6 | 1.1 | 7.7 |
