## Supplemental References for "Genomewide association study reveals transient loci underlying the genetic architecture of biomass accumulation under cold stress in Sorghum"

**SUPPLEMENTAL INFORMATION - REFERENCES**

72. Zhang, Z., Li, J., Pan, Y., Li, J., Shi, H., Zeng, Y., ... & Sun, X. (2017). Natural variation in CTB4a enhances rice adaptation to cold habitats. *Nature communications*, *8*, 14788.

106. Rui, Y., Xiao, C., Yi, H., Kandemir, B., Wang, J. Z., Puri, V. M., Anderson, C. T. (2017) POLYGALACTURONASE INVOLVED IN EXPANSION3 Functions in Seedling Development, Rosette Growth, and Stomatal Dynamics in Arabidopsis thaliana. *The Plant Cell Online*, tpc-00568.

122. Ye, W., Shen, C. H., Lin, Y., Chen, P. J., Xu, X., Oelmüller, R., Yeh, K., Lai, Z. (2014) Growth promotion-related miRNAs in Oncidium orchid roots colonized by the endophytic fungus Piriformospora indica. *PLoS One*, *9*(1): e84920.

139. Li, X., Yang, D., Sun, L., Li, Q., Mao, B., He, Z. (2016) The systemic acquired resistance regulator OsNPR1 Attenuates Growth by Repressing Auxin Signaling and promoting IAA-amido synthase expression. *Plant Physiology*, (2016): pp-00129.

140. Huda, K. M. K., Banu, M. S. A., Yadav, S., Sahoo, R. K., Tuteja, R., Tuteja, N. (2014) Salinity and drought tolerant OsACA6 enhances cold tolerance in transgenic tobacco by interacting with stress-inducible proteins. *Plant Physiology and Biochemistry*, 82: 229-238.

141. Zhang, G., Wang, F., Li, J., Ding, Q., Zhang, Y., Li, H., Zhang, J., Gao, J. (2015) Genome-wide identification and analysis of the VQ motif-containing protein family in Chinese cabbage (*Brassica rapa* L. ssp. *Pekinensis*). *International Journal of Molecular Sciences*, 16(12): 28683-28704.

142. Jeon, J., Kim, J. (2012). Arabidopsis response regulator 1 (ARR1) and Arabidopsis histidine phosphotransfer protein 2 (AHP2), AHP3, and AHP5 function in cold signaling. *Plant Physiology*, (2012): pp-112.

151. Zhu, J., Zwiewka, M., Sovero, V., di Donato, M., Ge, P., Oehri, J., Aryal, B., Hao, P., Linnert, M., Burgardt, N., Lücke, C. (2016) TWISTED DWARF1 mediates the action of auxin transport inhibitors on actin cytoskeleton dynamics. *The Plant Cell Online*, tpc-00726.

152. Su, C. F., Wang, Y. C., Hsieh, T. H., Lu, C. A., Tseng, T. H., Yu, S. M. (2010) A novel MYBS3-dependent pathway confers cold tolerance in rice. *Plant Physiology*, 153(1): 145-158.
